# Expansion of information in scientific research papers

**DOI:** 10.1101/2022.05.06.490896

**Authors:** Malika Abdullaeva, John J. Bromfield, I. Martin Sheldon

## Abstract

Presenting information in papers allows readers to see the evidence for the research claims. The amount of information presented to readers is increasing in high impact factor scientific journals. The aim of the present study was to determine whether there was a similar expansion in the amount of information presented to readers in subject-specific journals. We examined 878 research papers that were published in the journals Biology of Reproduction and Reproduction during the first six months of 1989, 1999, 2009, and 2019. Although there were few differences between the journals, we found that between 1989 and 2019 the number of figures increased 1.5-fold, the number of figure panels increased 3.6-fold, and the number of display items increased 5.6-fold. Amongst the display items, the number of images per paper increased 10-fold, and the number of graphs per paper increased 3.7-fold. The median paper in 1989 was 8 pages long, contained 6 tables and/or figures, with 1 image and 4 graphs. In 2019 the median paper was 12 pages long, contained 7 tables and/or figures, with 13 images and 15 graphs. This expansion of information in subject-specific journals implies that authors, reviewers, and editors need to help readers digest complex biological messages without causing information overload.

**Lay summary:** We are living in an age of science and information. The amount of information presented in research papers has increased over time in the top science journals. Our research examined whether there has been a similar expansion in information in two influential subject-specific journals. We counted how much information was presented in 878 research papers across a 30-year period in the journals *Biology of Reproduction* and *Reproduction*. There were few differences between the two journals. But there was a striking increase in the information presented to readers in 2019 compared with 1989. The typical paper in 1989 was 8 pages long and contained 1 picture and 4 graphs. In 2019 the typical paper was 12 pages long and contained 13 pictures and 15 graphs. This expansion of information means that subject-specific journals must balance the presentation of complex biological messages with the risk of causing information overload.

## Introduction

Journals like *Nature, Science* and *Cell* are thought to have an overwhelming and disproportionate effect on research impact and career progression ^1, 2, 3^. These high impact factor journals publish multidisciplinary research papers reporting novel, major advances of broad importance for a diverse readership. Commensurate with the influence of these high impact factor journals, authors are presenting increasing amounts of information to support their claims, and to meet the expectations of their readers, reviewers, and editors. On the other hand, most science is published in subject-specific journals that are read by scientists interested in a particular research field. It is unclear if the trend for presenting readers with abundant information in high impact factor journals is reflected by an expansion in information in subject-specific journals.

The most accessible information for readers of journals is that published in each research paper. This information is provided to readers in the text of the paper, in tables, and in figures. Clear scientific writing is important to help readers understand the main messages presented in papers ^4^. However, scientists particularly use figures to present their data, communicate information, and visualize ideas ^5, 6, 7^. Figures typically display images of photographs, graphs, gels, blots, drawings, and omics analyses. Readers often examine the figures along with the title and abstract of papers prior to reading the main text. PubMed facilitates this approach by providing readers with direct access to figures by clicking on thumbnail images of each figure, which are shown below the abstract on each PubMed Identifier landing page ^8^.

The amount of information presented to readers is increasing in journals that have a high impact factor. For example, between 1984 and 2004, the number of figure panels in biological research papers more than doubled in *Nature* and increased four-fold in *Cell*^3^. However, presenting more information in a paper than the reader has the ability or time to assimilate leads to data overflow, reader fatigue, diminished understanding, and information overload ^2, 9^. The aim of the present study was to determine whether there was evidence for an expansion in the amount of information presented to readers in subject-specific journals.

## Methods

### Journals

To quantify the information presented to readers of subject-specific journals, we selected the journals *Biology of Reproduction* (ISSN: 0006-3363) and *Reproduction* (ISSN: 1470-1626; formerly *Journal of Reproduction and Fertility*). Both *Biology of Reproduction* and *Reproduction* are well-respected journals that are supported by academic societies and have published primary research about reproductive biology since the 1960s. The 2019 *Journal Citation Reports* impact factor (Clarivate Analytics, Boston, USA) was 3.3 for *Biology of Reproduction* and 3.2 for *Reproduction;* the median impact factor for the 29 journals in the subject category was 2.8.

### Papers

We downloaded and examined the PDF version of each research paper published in *Biology of Reproduction* and *Reproduction* during the first six months of 1989, 1999, 2009 and 2019. Review papers were excluded from the analysis. We quantified the information presented in each of the papers. but we excluded supplementary files because this additional information is not presented directly to the reader of the paper in the printed, HTML or PDF versions, and was not available in 1989 or 1999.

For each PDF we counted the number of pages, independently numbered tables, and independently numbered figures. We also counted the number of independently labelled figure panels (Fig. 1). Most labels are lower- or upper-case letters, or roman numerals, but in some cases other labels are used (for example, a’). A composite or single element figure that had no labeled panels was counted as a single panel. Each PDF was examined independently by two people, and discrepancies in counts were identified and resolved. More than 95% of discrepancies between the observers was in counting labelled figure panels when labels were small, had low visual contrast within images, or were partially obscured by images or text.

**Figure 1.**
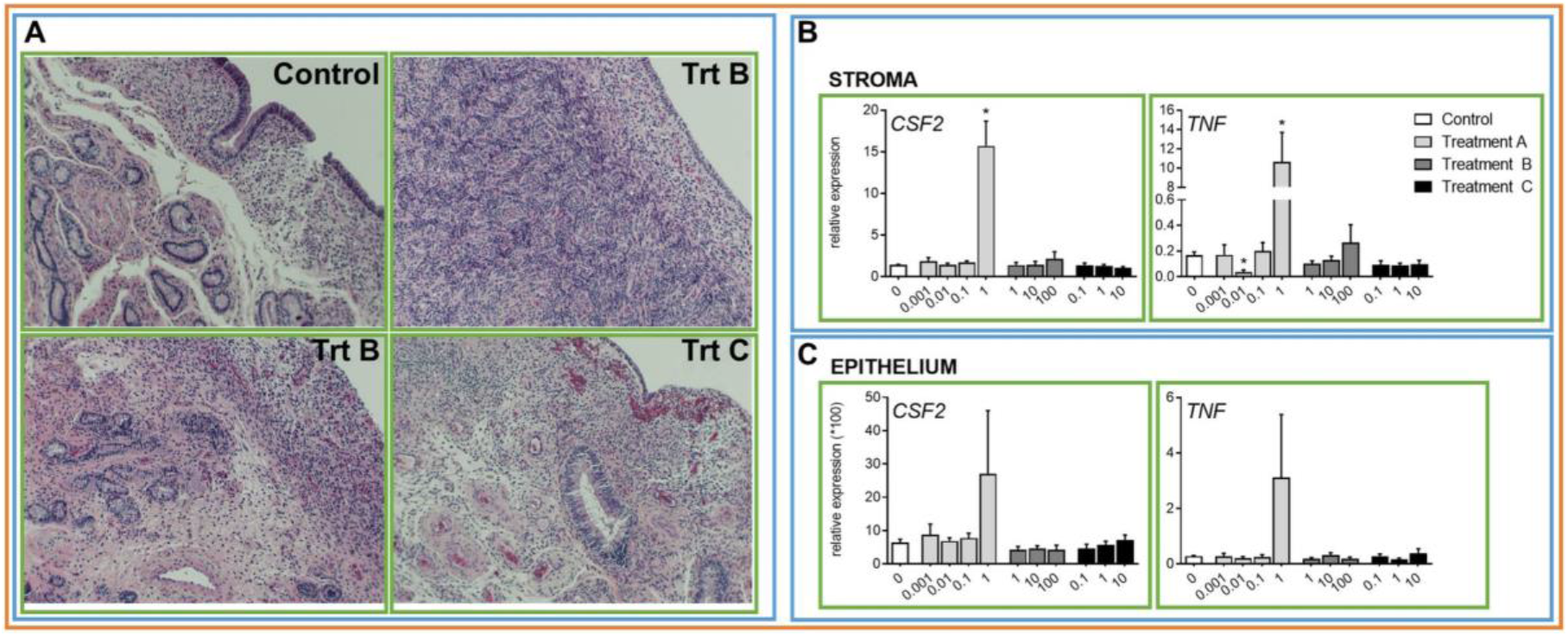
Figures, panels, and display items. A hypothetical example to illustrate the differences between an independently numbered figure (orange box), labelled figure panels (blue boxes), and display items (green boxes). This single figure contains three figure panels, and eight display items, of which four are images and four are graphs.

### Display items

Each independent element that comprises the whole or part of a figure can be called a display item (Fig. 1). Figures and even labelled figure panels often contain more than one display item. Display items were photographic images (such as photographs and photomicrographs), graphs (such as charts and flow cytometry plots), drawings (such as schematics and flow diagrams), images of gels (such as Western blots) and displays of outputs from omics analyses (such as heat maps and gene networks). Because the reader views each of these display items independently, we counted the number of each of these five types of display items in each paper. This also allowed us to calculate the total number of display items per paper.

### Statistical analysis

Data are reported as the arithmetic mean ± s.e.m., except where the median is reported (middle value in a sorted numerical list). Data were analyzed using SPSS version 28 (IBM Corp. Armonk, NY) and *p* < 0.05 was considered significant. Count data were analyzed for the factors of year, journal, and the interaction of year × journal. Statistical significance was determined using generalized linear models (GLiM) using negative binomial regression, and post hoc significance was demined using the Bonferroni adjustment for multiple comparisons ^10, 11^. Where there was overdispersion of count data due to a high proportion of zero values, statistical significance was also determined using Kruskal-Wallis tests for the factors of year and journal.

## Results

### Papers, authors, and pages

We examined 878 papers in *Biology of Reproduction* and *Reproduction* (Fig. 2A). The number of authors per paper did not differ significantly between the journals, but the number of authors per paper doubled between 1989 and 2019 (3.3 ± 0.1 vs. 7.2 ± 0.3, *p*= 10^−8^, Fig. 2B). There were also significant increases in the number of authors per paper between 1999 and 2009 (*p* = 0.001), and between 2009 and 2019 (*p* = 0.001).

**Figure 2.**
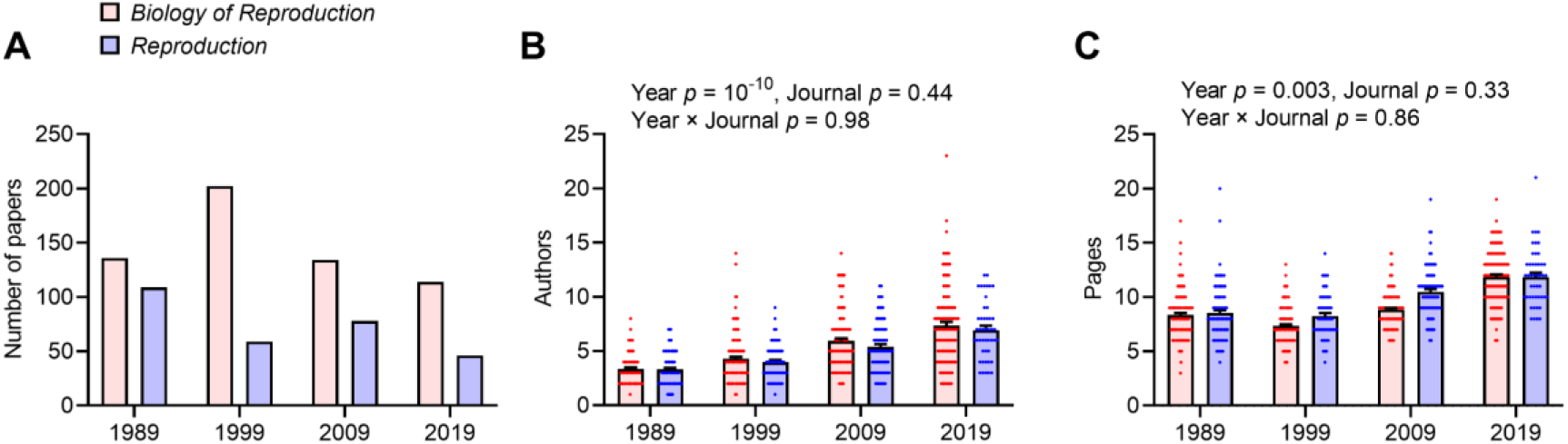
Papers, authors, and pages. (A) The number of research papers we examined that were published in *Biology of Reproduction* and *Reproduction* during the first six months of 1989, 1999, 2009 and 2019. (B) The number of authors per paper and (C) the number of pages per paper in *Biology of Reproduction* (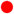, n = 586) and *Reproduction* (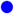, n = 292) during the first six months of 1989, 1999, 2009 and 2019. The width of distribution of points is proportionate to the number of points at that count, the bars represent the mean and the error bars the s.e.m.; statistical significance was determined by GLiM and *p*-values reported for each model.

The average number of pages per paper increased between 1989 and 2019 (8.4 ± 0.1 vs. 11.9 ± 0.1, Fig. 2C). The median number of pages per paper increased from 8 to 12 pages per paper between 1989 and 2019. The number of pages per paper did not differ significantly between the journals. The page size for *Biology of Reproduction* was 21.6 × 27.9 cm from 1989 to 2019, but the number of words per page increased from approximately 750 to 1100. The page size for *Reproduction* was smaller in 1989 (16.5 × 23.7 cm) and 1999 (19.6 × 26.6 cm) than in 2009 and 2019 (21.0 × 27.9 cm), and the number of words per page increased from approximately 700 in 1989, to 950 words per page in 1999, 2009 and 2019.

### Tables, figures, and figure panels

There was at least one table in 43.9% of papers and at least one figure in 97.3% of papers; only one paper did not contain either a table or a figure. The number of independently labeled tables did not differ significantly amongst the years, but there were more tables in *Reproduction* than *Biology of Reproduction* (Fig. 3A). Although there was no significant difference between the journals, the number of independently labeled figures increased 1.5-fold between 1989 and 2019 (Fig. 3B). The median for the total tables and figures per paper increased from 6 to 7 between 1989 and 2019 for both journals.

**Figure 3.**
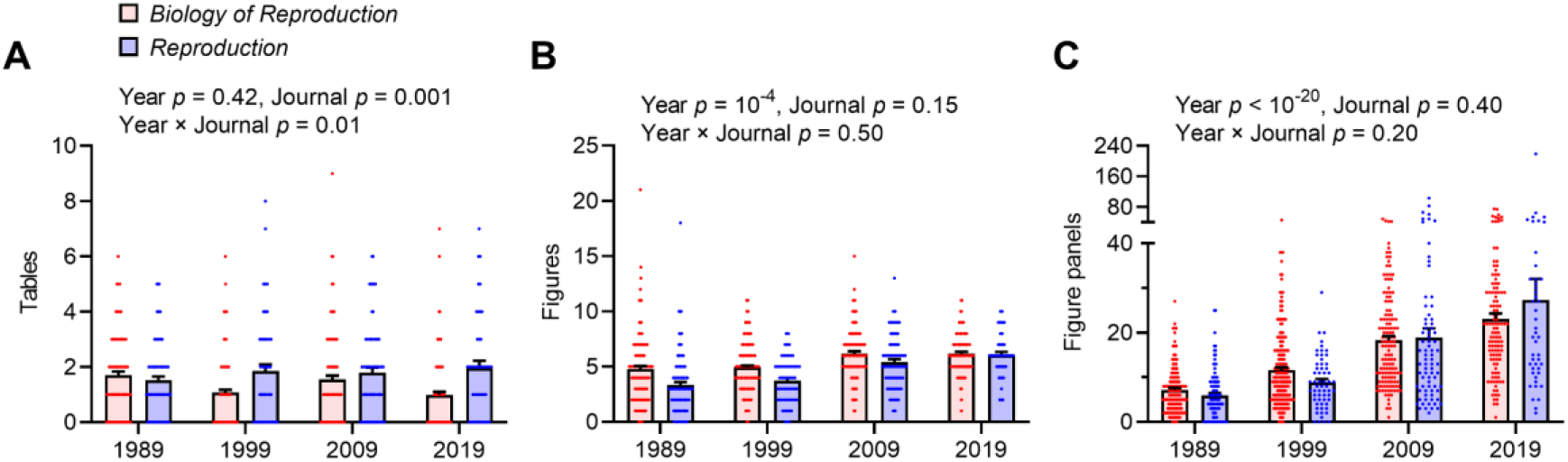
Tables, figures, and figure panels. The number of (A) independently numbered tables, (B) independently numbered figures, and (C) labelled figure panels, per research paper published in *Biology of Reproduction* (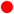, n = 586) and *Reproduction* (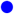, n = 292) during the first six months of 1989, 1999, 2009 and 2019. The width of distribution of points is proportionate to the number of points at that count, the bars represent the mean and the error bars the s.e.m.; statistical significance was determined by GLiM and *p*-values reported for each model.

As author instructions often limit the number of independently numbered figures, we also counted the number of labelled figure panels. Although there was no significant difference between the journals, there was a 3.7-fold increase in figure panels per paper between 1989 and 2019 (Fig. 3C). Similarly, the number of labelled figure panels per figure did not differ significantly between the journals but increased from 1.7 ± 0.2 in 1989 to 3.5 ± 0.9 in 2019 (*p* = 0.008).

### Display items in figures

There was a 5.6-fold increase in display items per paper between 1989 and 2019 (Fig. 4A). There was also a significant increase in display items per paper between each year (*p*< 0.001). There were minor differences in the number of display items per paper between the journals, but no significant differences between the journals within each year. The number of display items per figure did not differ significantly between journal (*p*= 0.9) but increased from 2.3 ± 0.3 in 1989 to 8.8 ± 1.9 in 2019 (*p*= 10^−6^). The number of display items per page also increased from 1989 to 2019 (1.0 ± 0.1 vs. 4.1 ± 0.2 display items per page, *p*< 0.001).

**Figure 4.**
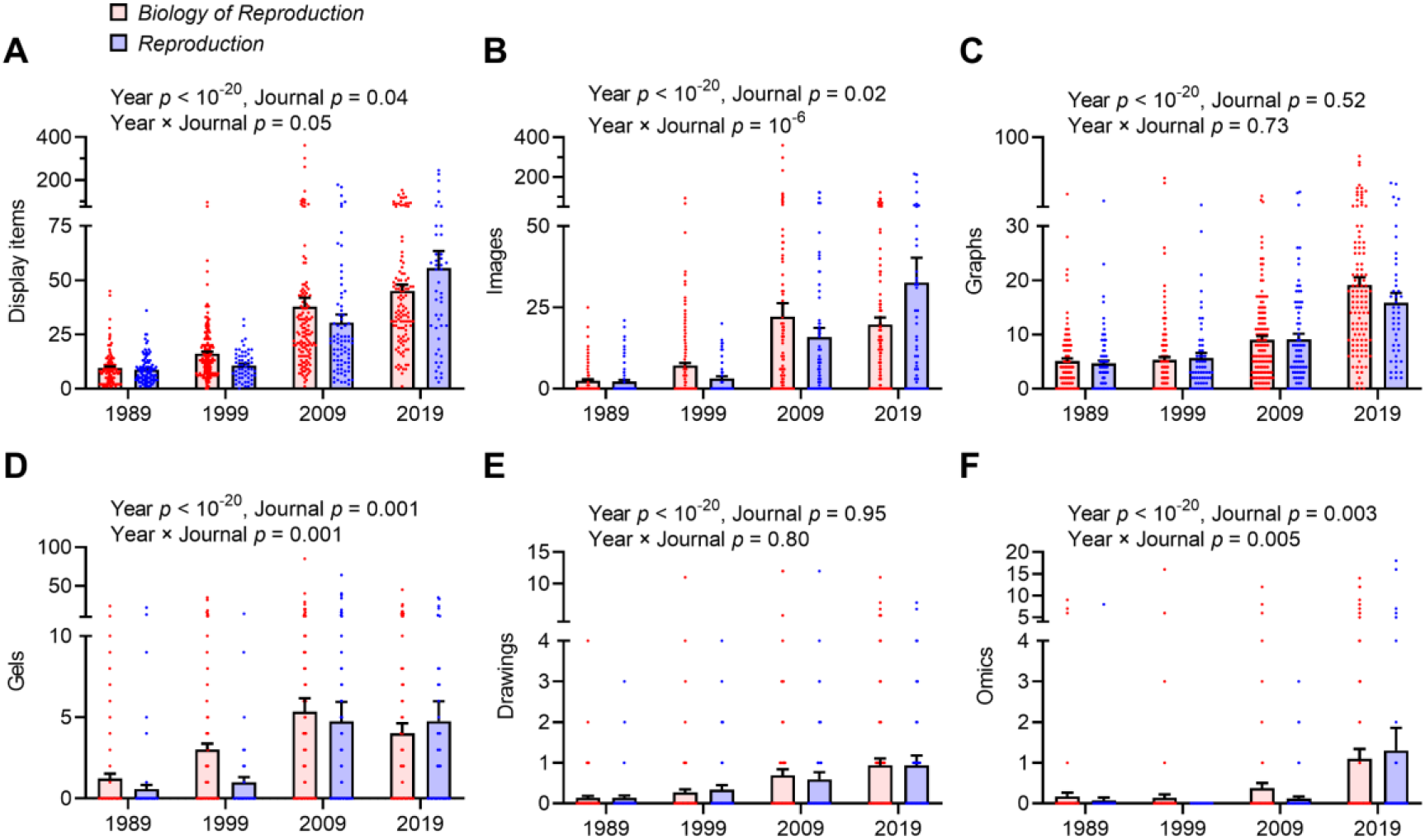
Display items in figures. (A) The number of display item per research paper published in *Biology of Reproduction* (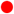, n = 586) and *Reproduction* (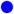, n = 292) during the first six months of 1989, 1999, 2009 and 2019. The display items include (B) images, (C) graphs, (D) gels, (E) drawings, and (F) representation of omics analyses, and the number of these items per research paper is presented. The width of distribution of points is proportionate to the number of points at that count, the bars represent the mean and the error bars the s.e.m.; statistical significance was determined by GLiM and *p*-values reported for each model.

Amongst the five types of display items, the most common were images and graphs. We found that 11.7% of papers displayed images, 42.1% of papers displayed graphs, and 42.1%of papers displayed both images and graphs. There was a 10-fold increase in the number of images per paper between 1989 and 2019 (Fig. 4B), and a 3.7-fold increase in the number of graphs per paper (Fig. 4C). Between 1989 and 2019, the median number of images increased from 1 to 13 and the median number of graphs increased from 4 to 15 per paper. There were more images per paper in *Biology of Reproduction* than in *Reproduction*, but the size effect differed within each year (Fig. 4B). There was no significant difference in the number of graphs per paper between the journals (Fig. 4C).

The number of gels, drawings, and presentations of omics analyses per paper also increased between 1989 and 2019 (*p* < 0.001, Fig. 4D-F). Although journal and the interaction of year × journal was significant for the number of gels and omics items in papers, the effect size was modest. Furthermore, the high percentage of counts with a zero value (gels 62.4%; drawings 78.4%, omics 92.1%) warrants caution in interpreting these statistical models ^11^. However, using Kruskal-Wallis tests as an alternative statistical approach for count data, we found similar statistical significance for year (gels *p*= 10^−20^, drawings *p*= 10^−20^, and omics analyses *p* < 10^−14^), and journal (gels *p*= 10^−5^, drawings *p*= 0.40 and omics analyses *p*= 0.04).

## Discussion

We found evidence for increasing amounts of information presented to the readers of the two subject-specific journals that we examined across a 30-year period. The expansion in information between 1989 and 2019 was associated with more figures, figure panels, and display items. The median paper in 1989 was 8 pages long, contained 6 tables and/or figures, with 1 image and 4 graphs, while in 2019 the median paper was 12 pages long, contained 7 tables and/or figures, with 13 images and 15 graphs. The increase in information presented in *Biology of Reproduction* and *Reproduction* papers since 1989 mirrors similar increases for biological research papers in *Nature* and *Cell* between 1984 and 2004 ^3^. The number of data items and panels per figure also doubled between 1993 and 2013 in a survey of 1,464 biological research papers across a selection of subject-specific and high impact factor journals^12^.

The benefits of presenting abundant information in papers include greater evidence for the research claims, better exclusion of alternative hypotheses, and a more comprehensive story. Authors can present more data now than 30 years ago because they have more research techniques, more equipment, more assays, and can make more measurements. Furthermore, there are easier workflows for collecting and displaying data such as transcriptomics, Western blots, and photomicrographs. Authors also have more powerful personal computers and software, making it easier to process images, draw graphs, perform statistical analyses, and assemble multipaneled figures. Digital workflows and on-line publication make it easier and cheaper to publish more data in papers or in supplementary files.

We found more figures, more figure panels, and more display items in 2019 than 1989 in subject-specific research papers. We suggest that the number of display items per paper was most useful for comparing information between papers. Counting each display item was straightforward and avoided potential confusion caused by unclear labelling of figure panels in some papers. Visualization of data helps researchers formulate their ideas as well as communicate information and concepts to the reader ^5^. Figures also engage the visual system and are often attractive to readers. The 5.6-fold increase in display items per paper since 1989, presents the reader of a 2019 paper not only with more display items to assimilate, but also longer figure legends. Figure legends typically contain technical details, symbols, and abbreviations. Authors need to craft these figure legends so that readers can clearly grasp the take-home messages ^13^.

Whilst it might be argued that more data were reported in the text in 1989 than 2019, we found that papers in 1989 were typically 4 pages shorter than papers in 2019. We also found that the number of tables did not differ significantly among the years, which may reflect the role of tables in presenting numbers and words in columns and rows that have to be read like text. The common statement of “data not shown” 30 years ago has been replaced by display items in the paper or supplementary files. In high impact factor journals, these supplementary files often contain more data than the paper ^3, 14^. Open access to research data in papers is associated with more citations, media attention, potential collaborators, job opportunities and funding opportunities ^15^. Open access to data in repositories is also desirable for specialist readers and facilitates secondary use of the data ^16^. Interestingly, papers that included links to data in a repository were predicted to have a 25% higher citation impact ^17^.

A benefit of peer-review is that reviewer suggestions often improve papers. Indeed, the greatest risk to the quality of published papers is indifferent reviewers accepting low-quality manuscripts ^18^. However, opinion leaders have expressed concerns that some reviewers can be overdemanding in their requests for more information ^3, 19^. Concerns include reviewers requesting unnecessary control experiments, asking for experiments that extend beyond a paper’s story, or seeking information that exceeds the scope of subject-specific journals ^20, 21^. Editors are also under pressure to maintain their journal reputation and impact factor, and to compete with other journals for the best papers in their subject. Editors perceive that publishing information-rich papers will help their impact factor, reputation, and media coverage ^3, 15^. However, to avoid information overload for readers, the reviewers and editors may need to scale their expectations between high impact and subject-specific journals.

Providing more information in papers has other unintended consequences, including lengthening the research and publication process for each project ^3^. In addition, we found that the number of authors per paper doubled between 1989 and 2019 in the journals. Furthermore, the ease of collecting data may result in less thought put into designing experiments and analyzing results. Inappropriate data presentation and flawed statistical analysis are particular problems ^22^. In a survey of 580 papers, less than 1 in 6 papers met all the good practice criteria for image-based figures ^6^. Problems included missing scale bars, misplaced or poorly marked insets, images or labels that were not accessible to colour-blind readers, and insufficient explanations of labels or components of images. The aim is to present data simply, clearly, and honestly without clutter, confusion, or deception so that readers can focus on the research story ^6, 23^. Fortunately, software, training, and advice on how to prepare display items is readily available ^5, 6, 7, 24, 25^.

The increasing information presented in research papers may also be influenced by human nature because we tend to solve problems by adding rather than subtracting components ^26^. Instead, simplifying the information presented in research papers, whilst other data are provided elsewhere, adds value for readers and reduces the risk of information overload. Simplification does not mean cherry-picking data or dumbing down information but making the story more comprehensible for readers. As authors, we should present the information in the paper that are needed to inform the story but confine other data to supplementary files or a data repository. Judicious use of graphical abstracts, careful structuring of the results section, and stylish writing also help. As reviewers, we need sufficient time, engagement, and knowledge to evaluate papers, so that we can suggest improvements that clarify the story. As editors, we should integrate the referee reports and communicate a consensus to the authors about the additional data and revisions required for their paper to be acceptable, and whether data presentation could be simplified without compromising the story.

In conclusion, we found more information presented to readers of research papers in two subject-specific journals over a 30-year period. Most striking was a 5.6-fold increase in display items per paper between 1989 and 2019. Providing abundant information generates a comprehensive scientific story and allows the readers to see the evidence for the research claims. However, the supply and demand for more information risks information overload for readers. Simplifying data presentation reduces the risk of information overload and helps readers digest complex biological messages in scientific research papers.

## Author contributions

Conceptualization, writing – original draft preparation, and supervision IMS; methodology, and formal analysis IMS, MA; visualization and writing – review and editing IMS, MA, JJB.

## References

1. Young NS, Ioannidis JPA, Al-Ubaydli O. Why current publication practices may distort science. PLOSMedicine 5, e201 (2008).

2. Siebert S, Machesky LM, Insall RH. Overflow in science and its implications for trust. eLife 4, e10825 (2015).

3. Vale RD. Accelerating scientific publication in biology. Proc Natl Acad Sci U S A 112, 13439–13446 (2015).

4. Woodford FP. Sounder thinking through clearer writing. A graduate course on scientific writing can, if appropriately designed, strengthen scientific thinking. Science 156, 743–745 (1967).

5. McGill GG. Knowledge synthesis through scientific visualization. Nature Microbiology 7, 185 (2022).

6. Jambor H, et al. Creating clear and informative image-based figures for scientific publications. PLOS Biology 19, e3001161 (2021).

7. Boers M. Designing effective graphs to get your message across. Ann Rheum Dis 77, 833–839 (2018).

8. Sayers EW, et al. Database resources of the national center for biotechnology information. Nucleic Acids Research 50, D20–D26 (2021).

9. Lanham RA. The economics of attention: Style and substance in the age of information. University of Chicago Press (2006).

10. Nelder JA, Wedderburn RWM. Generalized linear models. Journal of the Royal Statistical Society Series A (General) 135, 370–384 (1972).

11. Berk R, MacDonald JM. Overdispersion and poisson regression. Journal of Quantitative Criminology 24, 269–284 (2008).

12. Cordero RJB, de León-Rodriguez CM, Alvarado-Torres JK, Rodriguez AR, Casadevall A. Life science’s average publishable unit (APU) has increased over the past two decades. PLOS ONE 11, e0156983 (2016).

13. Kroodsma DE. A Quick Fix for Figure Legends and Table Headings. The Auk 117, 1081–1083 (2000).

14. Borowski C. Enough is enough. Journal of Experimental Medicine 208, 1337 (2011).

15. McKiernan EC, et al. How open science helps researchers succeed. eLife 5, e16800 (2016).

16. Wilkinson MD, et al. The FAIR Guiding Principles for scientific data management and stewardship. Scientific Data 3, 160018 (2016).

17. Colavizza G, Hrynaszkiewicz I, Staden I, Whitaker K, McGillivray B. The citation advantage of linking publications to research data. PLOS ONE 15, e0230416 (2020).

18. D’Andrea R, O’Dwyer JP. Can editors save peer review from peer reviewers? PLOS ONE 12, e0186111 (2017).

19. Ploegh H. End the wasteful tyranny of reviewer experiments. Nature 472, 391 (2011).

20. Raff M, Johnson A, Walter P. Painful publishing. Science 321, 36 (2008).

21. Walbot V. Are we training pit bulls to review our manuscripts? Journal of Biology 8, 24 (2009).

22. Vaux DL. Know when your numbers are significant. Nature 492, 180–181 (2012).

23. Cabanski C, Gilbert H, Mosesova S. Can graphics tell lies? A tutorial on how to visualize your data. Clinical And Translational Science 11, 371–377 (2018).

24. Wong B. Design of data figures. Nature Methods 7, 665 (2010).

25. Midway SR. Principles of Effective Data Visualization. Patterns (NY) 1, 100141 (2020).

26. Adams GS, Converse BA, Hales AH, Klotz LE. People systematically overlook subtractive changes. Nature 592, 258–261 (2021).

